# Host microRNA interactions with the SARS-CoV-2 viral genome 3’-untranslated region

**DOI:** 10.1101/2023.05.18.541401

**Authors:** Caleb J. Frye, Caylee L. Cunningham, Mihaela Rita Mihailescu

## Abstract

The 2019 pandemic, caused by the severe acute respiratory syndrome coronavirus 2 (SARS-CoV-2), has marked the spread of a novel human coronavirus. While the viral life cycle is well understood, most of the interactions at the virus-host interface remain elusive. Furthermore, the molecular mechanisms behind disease severity and immune evasion are still largely unknown. Conserved elements of the viral genome such as secondary structures within the 5’- and 3’-untranslated regions (UTRs) serve as attractive targets of interest and could prove crucial in furthering our understanding of virus-host interactions. It has been proposed that microRNA (miR) interactions with viral components could be used by both the virus and host for their own benefit. Analysis of the SARS-CoV-2 viral genome 3’-UTR has revealed the potential for host cellular miR binding sites, providing sites for specific interactions with the virus. In this study, we demonstrate that the SARS-CoV-2 genome 3’-UTR binds the host cellular miRNAs miR-760-3p, miR-34a-5p, and miR-34b-5p, which have been shown to influence translation of interleukin-6 (IL-6), the IL-6 receptor (IL-6R), as well as progranulin (PGRN), respectively, proteins that have roles in the host immune response and inflammatory pathways. Furthermore, recent work suggests the potential of miR-34a-5p and miR-34b-5p to target and inhibit translation of viral proteins. Native gel electrophoresis and steady-state fluorescence spectroscopy were utilized to characterize the binding of these miRs to their predicted sites within the SARS-CoV-2 genome 3’-UTR. Additionally, we investigated 2’-fluoro-D-arabinonucleic acid (FANA) analogs of these miRNAs as competitive binding inhibitors for these miR binding interactions. The mechanisms detailed in this study have the potential to drive the development of antiviral treatments for SARS-CoV-2 infection, and provide a potential molecular basis for cytokine release syndrome and immune evasion which could implicate the host-virus interface.

**Author Summary:** Severe acute respiratory syndrome coronavirus 2 (SARS-CoV-2) has now plagued the world for over three years. In this time, scientific advancements have allowed for the development of mRNA vaccines and targeted antiviral drugs. However, many mechanisms of the viral life cycle, as well as the interactions at the host-virus interface, remain unknown. The host immune response is of particular interest in combating SARS-CoV-2 infection, with observed dysregulation in both severe and mild cases of infection. To uncover the link between SARS-CoV-2 infection and observed immune dysregulation, we investigated host microRNAs associated with the immune response, particularly miR-760-3p, miR-34a-5p, and miR-34b-5p and emphasize them as targets of binding by the viral genome 3’-UTR. We utilized biophysical methods to characterize the interactions between these miRs and the SARS-CoV-2 viral genome 3’-UTR. Lastly, we introduce 2’-fluoro-D-arabinonucleic acid analogs of these microRNAs as disruptors of the binding interactions, with intent of therapeutic intervention.

## Introduction

SARS-CoV-2, the virus responsible for the COVID-19 pandemic, has caused nearly 6.9 million deaths as of May 2023.(1) Development of the COVID-19 mRNA vaccines has mitigated these numbers; however, the virus still poses a threat to those who are immunocompromised and/or have underlying health issues.(2) Additional treatment options do exist, however, the molecular mechanisms behind disease severity and the viral life cycle remain largely unknown, highlighting a need for elucidation of viral functions. Targeting the structural elements of the virus, including its genome, could serve as a prolific target and broaden our understanding of the viral life cycle. The SARS-CoV-2 viral genome is 29,903 nucleotides (nt) and harbors both 5’- and 3’-untranslated regions (UTRs), which have been shown to play a key role in viral replication through a plethora of conserved secondary structural elements, such as the stem-loop type 2 motif (s2m). (3–5) The s2m has been shown to act as an initiation site for dimerization through kissing interactions that can be converted to a thermodynamically favored extended duplex structure. Furthermore, this motif was also shown to bind host cellular microRNA (miR)-1307-3p *in vitro*. (6,7) With this in mind, we investigated whether the SARS-CoV-2 viral genome 3’-UTR could undergo additional interactions with host miRs, based on secondary structural elements.

Viruses are known to hijacking host elements to promote viral fitness, including miRs. SARS-CoV-1 and other viruses have been shown to alter miR expression, specifically of those that regulate immune cell proliferation and activation, as miRs provide a mechanistic route for direct effects on host translational regulation. (8,9) Computational analysis of the SARS-CoV genome revealed hundreds of potential miR-RNA binding interactions throughout the viral genome, centered around conserved secondary structural elements and the viral genome 3’-UTR. (10) Considering the high sequence homology and conservation of genomic structural elements between SARS-CoV and SARS-CoV-2, binding sites for host cellular miRs are predicted to be conserved as well, suggesting that SARS-CoV-2 may utilize similar interactions to aid in replication, transmission, and overall infectivity through loss of host translational regulation.(11,12) Thus, we and others analyzed the potential of the SARS-CoV-2 genome to bind to miRs which are implicated in host immune functions, particularly those related to the dysregulation of interleukin (IL) expression in various pathological conditions.(13–22) MiRs have been found to regulate the expression of specific ILs directly, or through regulatory signaling pathways.(13,14,23–25) In prostate cancer, host miR-760-3p has been shown to influence IL-6 providing a direct translational inhibitory function, as well as angiotensin converting enzyme 2 (ACE2) receptor, which is known as the primary host receptor targeted by SARS-CoV-2 for viral entry.(26–29) Additionally, miR-34a-5p was shown to decrease the expression of IL-6 and the IL-6 receptor (IL-6R) indirectly through plasminogen activator inhibitor-1 (PAI-1) in the JAK/STAT signaling pathway.(20,30,31) MiR-34b-5p has been identified as a direct regulator of PGRN, a pro-inflammatory protein known to activate cytokine production.(21,32) Recent work has also identified that both miR-34a-5p and miR-34b-5p could target the coding regions for both the viral membrane (M) and spike (S) proteins, respectively, which could allow for direct targeting of viral protein translation by the host immune response.(33) MiR-760-3p, miR-34a-5p, and miR-34b-5p all exhibit altered expression profiles upon SARS-CoV-2 infection, varying from slight reductions to a 2.94-fold decrease in serum levels. (9,19) The proposed roles of these miRs in host immunity has led to speculation that downregulation of these miRs may be favorable for SARS-CoV-2. (34)

Severe cases of COVID-19 are associated with the hallmark symptoms of cytokine release syndrome (CRS), also known as the “cytokine storm.” CRS is defined by a large increase in the expression of cytokines and cycles of inflammation that are capable of damaging nearby tissues.(35) Typically associated with a large virus titer, initiation of CRS is attributed to a large increase in the expression of several key cytokines, including IL-6 and IL-6R.(36) The pleiotropic nature of these cytokines leads to activation of several branches of the immune system across local tissues, inducing a spike in local immune cell activity and cycles of inflammation in the alveolar sacs, ultimately increasing the risk of respiratory failure and death. (35–39) Clinical data has also indicated that pro-inflammatory biomolecules like PGRN are overexpressed in the serum of patients with severe COVID-19, and it is believed that this overexpression contributes to the mass overexpression of ILs through a positive feedback loop.(40) Cycles of uncontrolled inflammation, triggered by the cooperative activation of both IL-6 and PGRN, are considered to be major factors in the initiation and progression of the cytokine storm in severe COVID-19 and can be linked to a loss of translational regulation.(35–38,41)

Considering the role of miR-760-3p, miR-34a-5p, and miR-34b-5p in the regulation of IL-6, IL-6R, and PGRN, these miRs serve as potent cellular targets for viral hijacking by SARS-CoV-2. With this in mind, we investigated whether SARS-CoV-2 could bind host cellular miR-760-3p, miR-34a-5p, and miR-34b-5p, as interactions with these miRs could allow for the virus to alter the host immune response, potentially benefitting the viral life cycle and contributing to greater viral fitness. Furthermore, intervention by antisense oligonucleotides could serve as therapeutic options to prevent viral hijacking of these miRs. Antisense oligonucleotides, like 2’-fluoro-D-arabinonucleic acids (FANAs), have been employed in several applications as binding inhibitors or in siRNA knockdowns, and have been theorized and developed to be effective oligos due to their ability to self-deliver *in vivo* and prove effective in many *in vitro* systems. (42–44) Often, antisense oligonucleotides have been employed for these purposes, designed as sequence specific targets, such as for siRNA-mediated knockouts or targeted binding. (42–45) Thus, FANAs could sequence-specific target these binding interactions and allow for a potential mechanism of intervention. In this work, we show that the SARS-CoV-2 viral genome 3’-UTR binds host cellular miR-760-3p, miR-34a-5p, and miR-34b-5p, and that these binding interactions can be inhibited by intervention with antisense FANA oligonucleotides complementary to the genome miR binding sites.

## Materials and Methods

### Oligonucleotide Synthesis

Selected sequences for the miR-760-3p, miR-34a-5p, and miR-34b-5p binding sites were based on the predicted fold of the SARS-CoV-2 genome 3’-UTR, as determined by the software program RNAstructure.(46) The RNA and FANA sequences used in this study (Table 1) were chemically synthesized and purchased from Dharmacon, Inc. and AUM Biosciences, LLC., respectively. The lyophilized samples were resuspended in sterile cacodylic acid (10 mM, pH 6.5) prior to use.

**Table 1:**
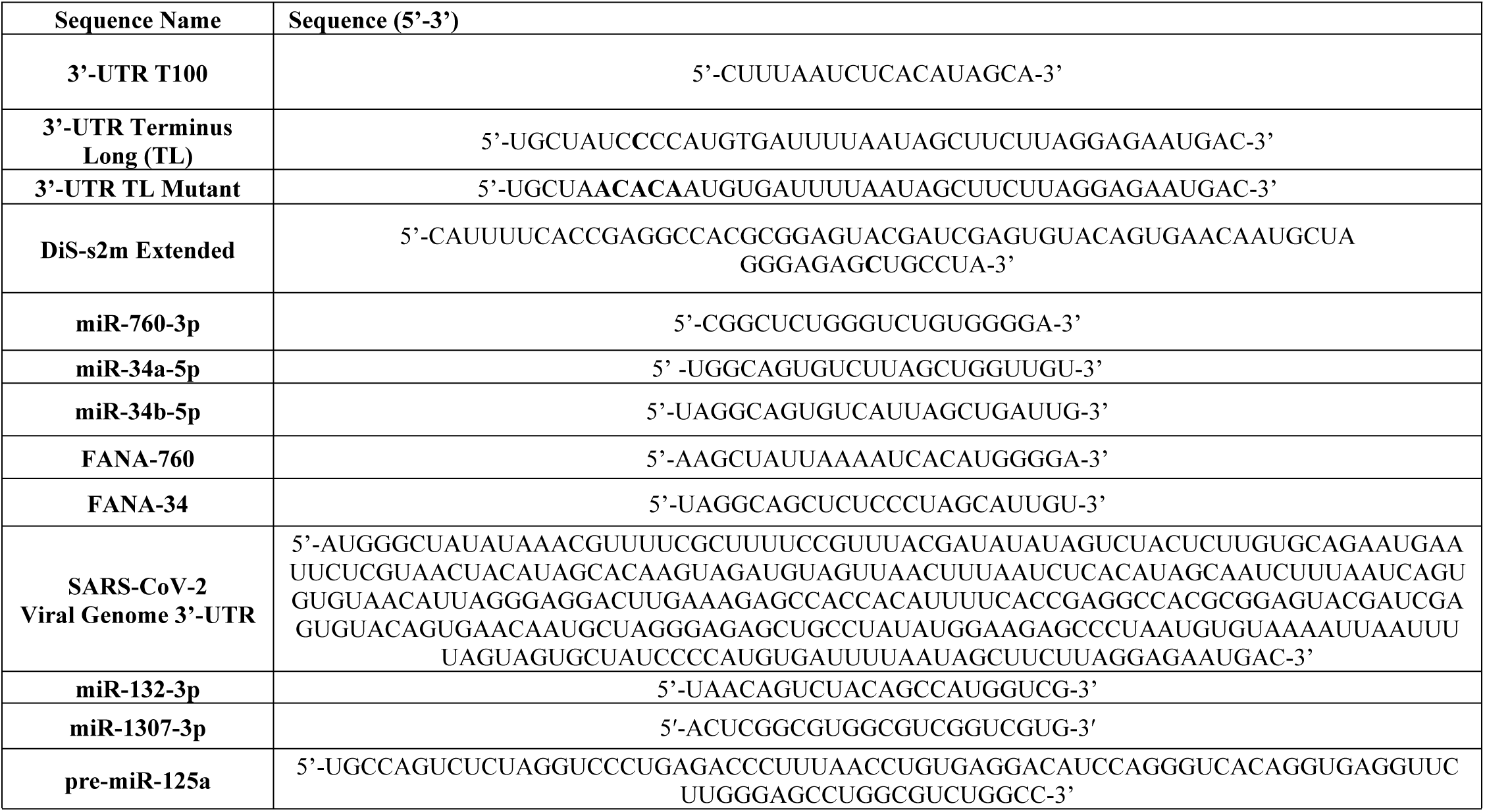
Selected oligonucleotides used in this study. All SARS-CoV-2 3’-UTR sequences were purchased from Dharmacon (Horizon Discovery Biosciences, PLC), and the FANA sequences were purchased from AUM Biosciences. The SARS-CoV-2 genome 3’-UTR was synthesized in-house (see methods).

The full-length SARS-CoV-2 genome 3’-UTR construct (a kind gift from Dr. Anna Wacker, Institute for Organic Chemistry and Chemical Biology, Germany) was transcribed from a psp64 dsDNA plasmid containing the sequence for the full-length 3’-UTR, a HINDIII recognition site, and a T7 RNA polymerase promoter upstream of the SARS-CoV-2 genome 3’-UTR sequence, as described previously.(5) The plasmids were transfected and amplified in DH5α *Escheria coli*, and extracted using a Qiagen Plasmid

GigaKit (Qiagen, Valencia, California, USA). The extracted plasmid DNA was reconstituted in cacodylic acid (10 mM, pH 6.5) and subjected to linearization by HINDIII-HF restriction digestion (New England Biolabs). Following digestion, the linearized plasmids were purified by three rounds of phenol:chloroform:isoamyl alcohol (PCI) phase extraction, lyophilized, and reconstituted in cacodylic acid (10 mM, pH 6.5). The purified plasmid DNA was used as a template for transcription by T7 RNA polymerase, and the reaction was terminated by addition of DNase I and proteinase K (New England Biolabs). To desalt the RNA while maintaining its native fold, the 3’-UTR RNA construct was run on a PD10 desalting column equilibrated with sterile cacodylic acid (10 mM, pH 6.5).

### Native Polyacrylamide Gel Electrophoresis

To test the binding of miR-760-3p to the SARS-CoV-2 3’-UTR, we utilized sequences mimicking the respective miR binding site located within the genome 3’-UTR. The T100 and TL sequences (Fig. 1, turquoise, Table 1) were prepared to pre-form a duplex which matches the native fold of the 3’-UTR at the miR-760-3p binding site. To promote formation of this duplex, the T100 and TL samples were brought to 100°C then allowed to slow anneal in a water bath for 1 hour until reaching room temperature (23°C). Two additional samples, the free T100 and TL sequences, were each boiled for 5 minutes and snap-cooled. 1 mM MgCl_2_ was added to a fixed 1 μM 3’-UTR T100:TL duplex mimic, along with increasing stoichiometric additions of miR-760-3p (which was previously snap-cooled): 0.5 μM, 1 μM, 1.5 μM, 2 μM. An additional control sample was prepared with all three sequences (T100, TL, and miR-760-3p) at 1 μM slow annealed together to promote the binding of miR-760-3p to the T100:TL duplex mimic. All samples were incubated on the lab bench at 23 °C for 1 hour, and loaded on 12% acrylamide:bisacrylamide tris-boric acid with 5 mM MgCl_2_ (TBM) gels which were run at 75V for 4 hours in ½x TBM buffer. Following electrophoresis, the TBM gel was stained in SYBR gold.(47) The gels were visualized using UV transillumination at 302 nm on a ProteinSimple AlphaImager HP. The negative binding control experiments utilizing miR-34a-5p and miR-1307-3p, along with the binding experiments with the TL mutant sequence, were repeated in the exact procedure as the described above for miR-760-3p PAGE experiments. Each control experiment with the respective RNAs was repeated in triplicate.

**Fig. 1:**
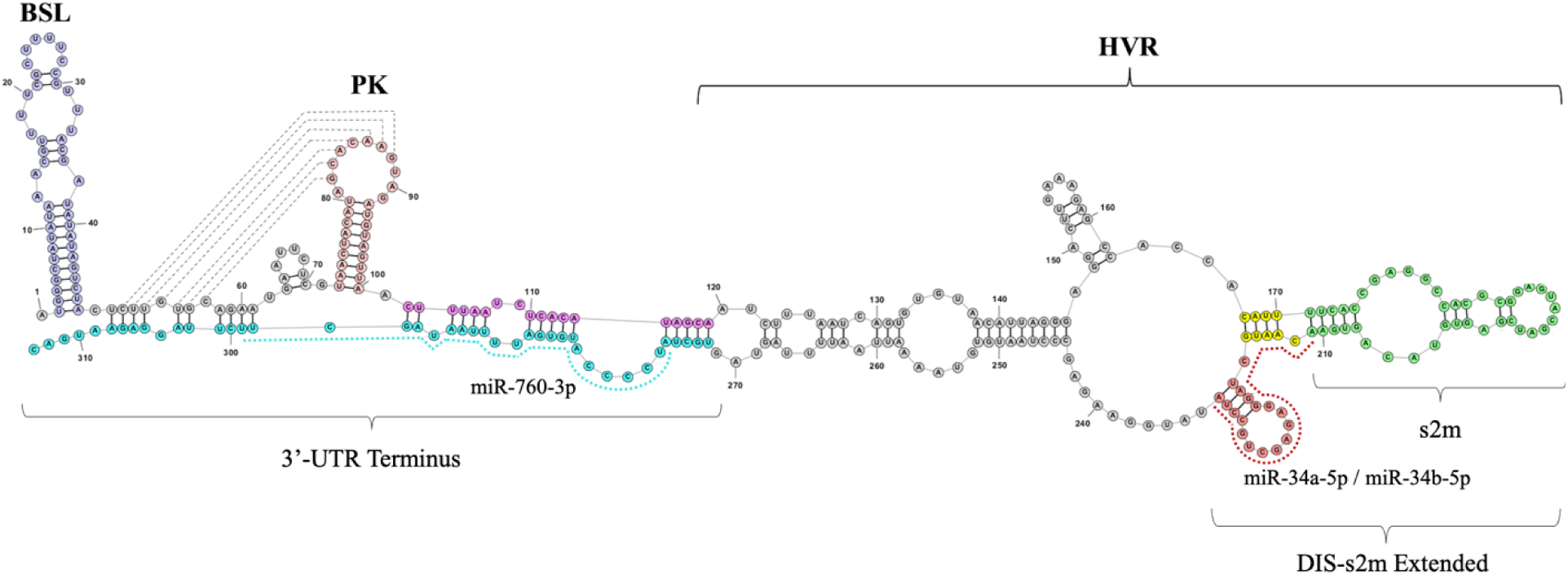
Schematic diagram of the SARS-CoV-2 genome 3’-UTR highlighting structural elements. The secondary structure of the 3’-UTR, as predicted by RNAstructure and StructureEditor software packages, was used as a structural basis for miRNA binding site predictions.^53^ Host cellular miR-760-3p has a binding site present at the 3’-UTR Terminus (turquoise, dotted line), and host cellular miRNAs miR-34a-5p and miR-34b-5p have a binding site downstream of the s2m element, which extends into the lower stem of the s2m (red, dotted line). Highlighted are also conserved structural elements like the bulged-stem-loop (BSL), the pseudoknot (PK), as well as the hypervariable region (HVR) containing the s2m element.

The miR-34a-5p and miR-34b-5p binding experiments utilized sequences which mimic the respective binding site for both miRs on the SARS-CoV-2 genome 3’-UTR. The sequence construct, named DiS-s2m extended (or s2m extended), was snap cooled on dry ice for 5 minutes, then brought to 23°C on the bench top. A snap-cooled stock of miR-34a-5p was added in increasing concentrations (0.25 μM, 0.5 μM, 0.75 μM, 1 μM, 1.5 μM, and 2 μM) in the presence of MgCl_2_ (1 mM), and the samples were incubated for one hour at room temperature (23°C), and electrophoresed on 15% TBM gels for at 75V and 4°C for 4 hours. The gels were then visualized using SYBR gold stain for identification of all RNA complexes. The experiment was repeated for miR-34b-5p, utilizing the same stoichiometric ratios and procedure. All experiments were performed in triplicate. An additional negative control for the DIS-s2m extended binding experiments was performed using miR-132-3p, which was previously used as a negative binding control for native PAGE experiments of the isolated s2m.(6) The binding experiments for miR-132-3p were performed in duplicate.

The fluorescently labeled miR-760-3p, miR-34a-5p, and miR-34b-5p were synthesized by Dharmacon (Horizon Discovery). The miR-760-3p sequence was labeled with the fluorophore DY547 (excitation: 558 nm, emission: 574 nm), and the miR-34a-5p and miR-34b-5p sequences were labelled with the fluorophore Cy3 (excitation: 554 nm, emission: 568 nm). All sequences were tagged at the 5’ end. The wild-type miR-760-3p, miR-34a-5p, and miR-34b-5p binding PAGE experiments were repeated with the fluorescently tagged miRs, and results were obtained in triplicate.

The FANA-760 and FANA-34 binding experiments were replicated using the same native PAGE procedure as the wild-type miRs. The electrophorized gels were stained in SYBR gold, and repeated in triplicate. All binding native PAGE binding experiments for both miRs and FANAs were done in triplicate.

### Steady-State Fluorescence Spectroscopy

Steady-state fluorescence spectroscopy experiments were performed at 25°C on a Horiba Jobin Yvon Fluoromax-4C with accompanying software, fitted with a 150 W ozone-free xenon arc lamp. The experiments were performed using a 3-mm path-length quartz cuvette (Starna cells) the samples being 150 μL. The excitation wavelength was set to 350 nm for pyrollo-cytosine (pyrC) and emission data was acquired from 400 to 500 nm.

The 3’-UTR T100:TL duplex mimic was prepared using a pyrC tagged Terminus Long sequence (pyrC-TL, pyrC is bolded in Table 1) and T100 sequence at 150 nM in 10 mM cacodylic acid, pH 6.5 which were slow annealed and incubated with 1 mM MgCl_2_ as described above for the miR-760-3p PAGE experiment. Snap-cooled miR-760-3p was titrated in 12.5 nM increments to the 3’-UTR T100:TL duplex mimic and equilibrated for 15 minutes prior to recording the emission intensity at 445 nm. Data points were corrected using blank subtraction and miR noise subtraction and normalized to the free 3’-UTR duplex emission intensity. These experiments were repeated in triplicate, and data was fit to equation 1 to determine the dissociation constant K_d_.

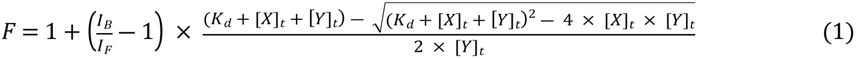

In equation 1, I_B_/I_F_ represents the ratio of fluorescent intensities of the bound and free RNA states (in this case pyrC TL), and [X]_t_ and [Y]_t_ represent the concentration of the titrant miR or full length 3’-UTR (in this case miR-760-3p), respectively. The K_d_ and respective error were reported as an average of the K_d_ from triplicate experimentation, and the error reported is the standard deviation of those values. The steady-state fluorescence experiments were repeated for the FANA-760 to determine the K_d_ of FANA-760 for the 3’-UTR T100:TL duplex mimic, and results were obtained in triplicate. The data was normalized and fit to equation 1, where Y represented in this case FANA-760.

For analysis of miR-34a-5p and miR-34b-5p binding to the DIS-s2m extended, similar experiments were performed using the same stoichiometric ratios of RNA as the miR-760-3p binding experiments and a pyrollo-cytosine labeled DIS-s2m extended (the substituted C is bolded in Table 1). The miR was added in 25 nM increments to a final concentration of 150 nM. These experiments were repeated in triplicate for each miR, and for the FANA-34. The collected data was normalized and fit to equation 1, in which case [X] represents either miR-34a-5p, miR-34b-5p or FANA-34 and [Y] represents the DIS-s2m extended, determining the respective K_d_ values for each complex.

For a negative binding control of fluorescence experiments to the 3’-UTR T100:TL duplex mimic, we used miR-34a-5p, whereas for a negative control of binding to the DIS-s2m extended we used miR-132-3p, with the binding curves for the negative controls being reported as an average of triplicate experiments.

To assess binding of the miR-760-3p, miR-34a-5p, and miR-34b-5p to the full length 3’-UTR, a modified version of the previously described experiments was utilized. The DY547 (miR-760-3p) and Cy3 (miR-34a-5p and miR-34b-5p) fluorescent-tagged miRs were prepared in cacodylic acid (10 mM, pH 6.5) at a final concentration of 150 nM RNA. For the DY547 fluorophore, excitation wavelength was set to 558 nm, and the emission was recorded at 574 nm. For the Cy3 fluorophore, the excitation wavelength was set to 550 and emission was recorded at 563 nm. The full length 3’-UTR was titrated in 25 nM increments to the tagged miRs, and the normalized binding curves for each miR were fit to equation 1, where in this case [X] is the 3’-UTR and [Y] is either of the DY547/Cy3-tagged miRs. K_d_ values and respective errors are reported for each miR as an average of triplicate fits. The K_d_ values of each miR to the full-length 3’-UTR were compared to the reported K_d_ binding affinities for the smaller constructs (3’-UTR terminus, DIS-s2m extended) using a two-tailed Student’s T-test with equal variances, to define statistical significance between binding to the small constructs and the full-length 3’-UTR. The Student’s T-test was performed on the raw data points for each binding curve, with comparisons drawn between the averages of each raw data set for each miR.

To establish specificity of the miR for the SARS-CoV-2 3’-UTR, a negative control was run using a construct of pre-miR-125a (86 nt, Horizon Discovery Biosciences, LTD) in place of the full-length 3’-UTR. The experiment was repeated in triplicate for each miR, and the reported binding curve was prepared by taking the average of each triplicate experiment for each miR. A representative curve was obtained by averaging the negative controls to represent all nonbinding experiments for the full-length 3’-UTR binding experiments.

To determine the efficiency of the FANA-760 and FANA-34 oligomers as binding inhibitors, we performed competition experiments to the full-length 3’-UTR between the FANAs and the fluorescent DY547/Cy3-tagged miRs. A sample of 150 nM DY547/Cy3-tagged miR was incubated with 250 nM of the 3’-UTR full length construct, followed by titration of FANA-760 or FANA-34 monitoring the increase in fluorescence intensity at 563 nm for Cy3 or 574 nm for DY547 as the respective FANA competed the miR. The normalized intensity was recorded and fitted to equation 2 for determination of the IC50 value (AAT Bioquest, equation adapted in Origin Pro) for the experiment.(48–52) In equation 2, F_max_ and F_min_ represent the maximum and minimum fluorescent intensities of the experiment. F_o_ represents the fluorescent intensity at each data point, and [S]_o_ represents the concentration of competing ligand, in this case the FANAs or untagged miRs. The K_I_ values for FANA binding were calculated using a modified Cheng-Prusoff equation (equation 3) based on the determined IC50, with [L] as the concentration of labeled miR and K_d_ being the experimentally derived K_d_ values for the labelled miR to the full-length 3’-UTR, as determined by equation 1.(53) The IC50 for each FANA was reported as an average of triplicates, and the reported errors were the standard deviation of the data set. For the K_I_, the reported values are the average of the calculated K_I_ from each IC50 value, and the error reported is the standard deviation of the K_I_ values. For comparison and evaluation of the determined IC50 and K_I_ values for the FANAs, the competition experiments were repeated, this time titrating unlabeled miR-760-3p, miR-34a-5p, or miR-34b-5p in place of FANA-760 or FANA-34. The IC50 values for the miRs were determined using equation 2, and compared to the IC50 values of the respective FANAs.

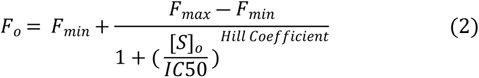

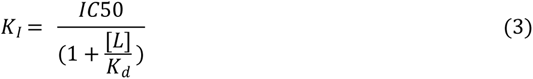

## Results

We and others have analyzed the SARS-CoV-2 3’-UTR for potential binding sites for host miRs (6,7,11,33,54,55) and identified that miR-760-3p has a predicted binding site located at nt 29,833 – 29,856 in the reference SARS-CoV-2 genome (NCBI: NC_045512.2) (Fig. 1, nt 272-313 in the 3’-UTR, turquoise dashed line) and that miR-34a-5p and miR-34b-5p both have the same predicted binding site, downstream of the s2m, extending into the lower stem of the s2m (Fig. 1, nt 211-233, nt 29,768 – 29,790 in the reference genome, red dashed line). Thus, we tested if these miRs bind to their predicted sites, both in model system comprised of short oligonucleotides that mimic the respective binding site, as well as in the context of the full length 3’-UTR.

### Host Cellular miR-760-3p Interacts with the SARS-CoV-2 Genome 3’-UTR Terminus

We first analyzed the binding of miR-760-3p to the SARS-CoV-2 genome 3’-UTR by native PAGE. In the context of the entire genome 3’-UTR, miR-760-3p is predicted to initiate binding at the exposed six-nucleotide bulge located at the 3’-UTR terminus; thus, to mimic this binding site, we pre-formed a duplex structure (Fig. 2A) using two chemically synthesized sequences (Table 1): first, from nt 102-119 in the full-length 3’-UTR, named here T100 (purple in Fig. 1 and 2A) and the second, from nt 272-313 of the full-length 3’-UTR, named here TL (turquoise in Fig. 1 and 2A).

**Fig. 2:**
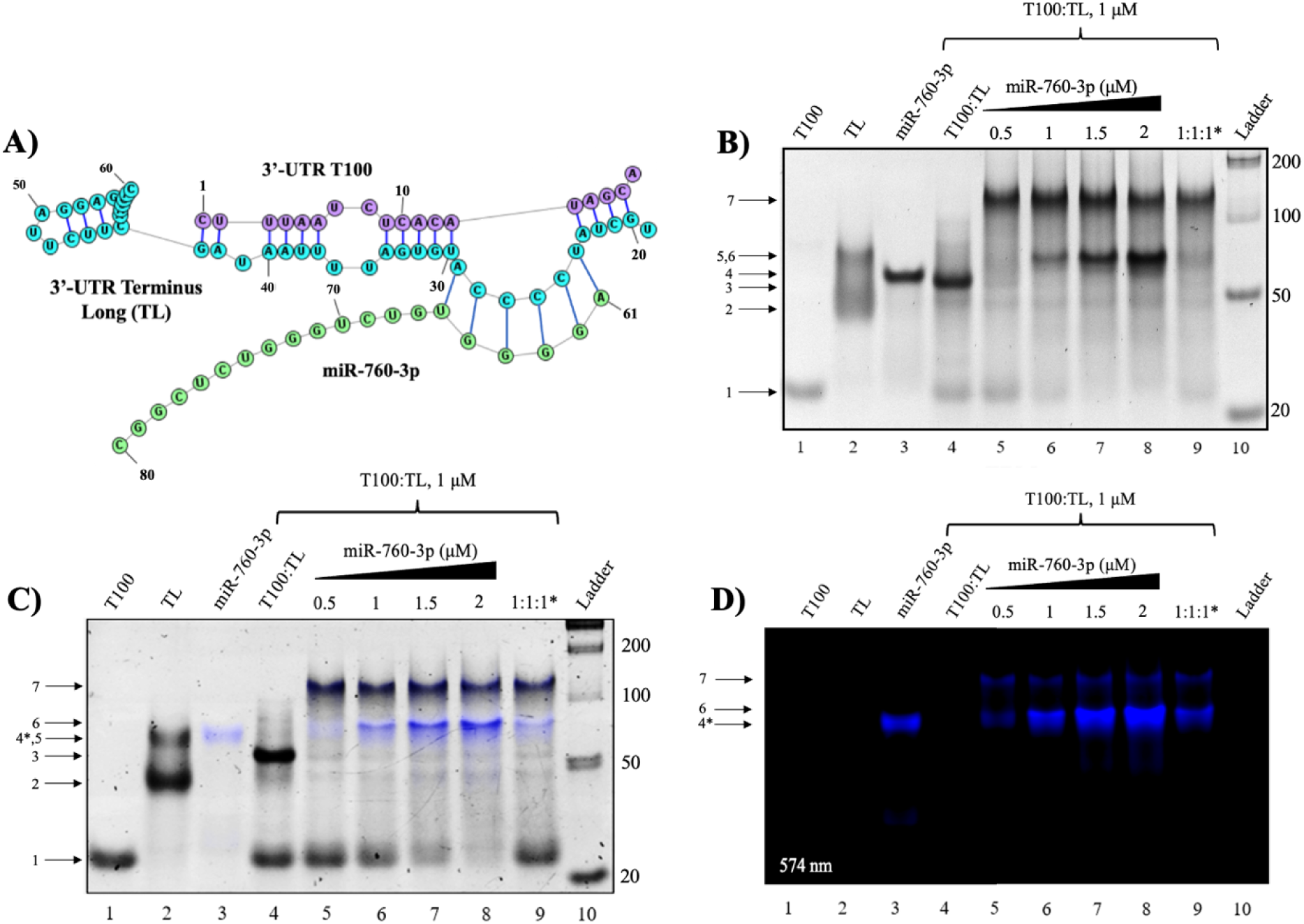
Native PAGE analysis of miR-760-3p binding to the 3’-UTR duplex mimic. **(A)** The predicted structure of miR-760-3p bound to the preformed T100:TL duplex mimic. **(B)** Native PAGE of the miR-760-3p binding to the 3’-UTR T100:TL duplex mimic. The free T100 sequence (lane 2, arrow 1) and free TL sequence (lane 2, arrows 2 and 5) were slow annealed to form the 3’-UTR duplex mimic (lane 4, arrow 3). MiR-760-3p (lane 3, arrow 4) was titrated to the 3’-UTR duplex mimic and two higher molecular weight complexes were identified. The lower weight band (lanes 5-9, arrow 6) is attributed to a complex of the 3’-UTR T100:Tl duplex mimic bound to one copy of miR-760-3p, and the higher molecular weight band (lanes 5-9, arrow 7) is attributed to a dimer of the 3’-UTR T100:TL duplex mimic bound to one copy of miR-760-3p. **(C)** Identical Native PAGE gel of the fluorescently tagged DY547-miR-760-3p binding to the 3’-UTR T100:TL duplex. All band identities are consistent with the original binding experiment; however, we observe altered migration of the DY547-miR-760-3p due to the DY547 positive charge (arrow 4*). **(D)** Original fluorescent image taken at 545 nm reveals the signature of the DY547-miR-760-3p (arrow 4*) indicating its presence in the two larger complexes that match the upper complex bands (arrows 6 and 7) observed in the original experiment.

In a TBM gel, in the presence of Mg^2+^, the isolated T100 migrates as a monomer (Fig. 2B, lane 1, arrow 1), the TL is present as a mixture of monomer and dimer (Fig. 2B, lane 2, arrows 2 and 5), and miR-760-3p forms a higher molecular weight complex that is likely a trimer (Fig. 2B, lane 3, arrow 4). When slow annealed together, TL forms a stable duplex with T100, as evidenced by the appearance of a new band (Fig. 2B, lane 4, arrow 3) corresponding to the complex’s predicted size of 60 nt. Upon the addition of increasing concentrations of miR-760-3p, this band disappears with the concomitant appearance of two new upper bands (Fig. 2B, arrows 6 and 7). One of these bands (Fig. 2B, arrow 6) is attributed to the formation of a complex between the 3’-UTR T100:TL duplex and one copy of miR-760-3p, with a predicted size of 80 nt (Fig. 2A). The second band (Fig. 2B, arrow 7) is attributed to a dimer of this observed complex, with a predicted size of 160 nt, as the 3’ extension (nt 43-60) of the 3’-UTR T100:TL duplex is predicted to be able to dimerize. To confirm that miR-760-3p is binding to the predicted exposed bulge, we mutated the TL sequence at that site (Fig. 2A, nt 24-29, bolded in Table 1), which is referred to as the 3’-UTR TL mutant. Repetition of the miR-760-3p binding experiments revealed that by mutating the exposed bulge on the 3’-UTR T100:TL duplex abolished the binding of miR-760-3p, confirming that this miR interacts with the 3’-UTR T100:TL duplex mimic at that site (S1 Fig. 1).

To establish the identity of miR-760-3p in the higher molecular complexes from previous native PAGE experiments, we labeled miR-760-3p with a DY547 fluorescent tag at its 5’ end (named here DY547-miR-760-3p), and repeated similar binding experiments (Fig. 2C and 2D). The DY547 fluorescent tag (excitation: 558 nm; emission: 574 nm) carries a positive charge, which leads to altered migration patterns of the DY547-miR-760-3p as compared with the unlabeled wild type miR-760-3p (Fig. 2C and 2D, lane 3, arrow 4*). Upon titration of the DY547-miR-760-3p to the 3’-UTR T100:TL duplex the two upper bands we previously assigned to the 3’-UTR T100:TL-miR-760-3p complex (arrow 6) and to its dimer (arrow 7) have the fluorescent signature of the DY547 tag as seen in an overlay of the fluorescent image and the SYBR gold stain (Fig. 2C, lanes 5-9, arrows 6 and 7). Thus, fluorescent PAGE analysis of DY547-miR-760-3p binding confirms the presence of miR-760-3p in both upper complexes (Fig. 2C and 2D, arrows 6 and 7). Further native PAGE experiments were performed to demonstrate that the predicted binding site for miR-760-3p is specific for this miR. MiR-34a-5p and miR-1307-3p were used as negative controls and, as expected, neither miR binds to the 3’-UTR T100:TL duplex (S1 Fig. 2). Taken together, these experiments show that host miR-760-3p binds to the SARS-CoV-2 genome 3’-UTR in its native fold, as we observe no disruption of the original 3’-UTR T100:TL duplex mimic after binding with miR-760-3p.

Quantitative information about miR-760-3p binding to the 3’-UTR T100:TL duplex was obtained by steady-state fluorescence spectroscopy. A modified 3’-UTR TL sequence containing a pyrollo-cytosine tag (pyrC, excitation: 350 nm; emission: 445 nm) in the exposed bulge of the miR-760-3p binding site (5’-UCpyrCCCA-3’, bolded in Table 1) was used in the preformation of the 3’-UTR T100:TL duplex mimic, followed by titration of the wild-type miR-760-3p. The binding curve (Fig. 3A) was fitted with equation 1 to determine the dissociation constant, K_d_. These experiments were performed in triplicate with the reported error being the standard deviation. We determined a K_d_ of 24.0 ± 4.1 nM (R^2^ = 0.9969) for the binding of miR-760-3p to the 3’-UTR T100:TL duplex mimic (Fig. 3A). Next, we assessed the binding of miR-760-3p to the full-length SARS-CoV-2 3’-UTR. In these experiments we used the fluorescently labeled DY547-miR-760-3p, monitoring the quenching of its fluorescence upon its binding to the unlabeled full-length 3’-UTR (Fig. 3B). Similarly, these experiments were performed in triplicate with the reported error being the standard deviation. For the complex formed by DY547-miR-760-3p with the full-length 3’-UTR, we determined a K_d_ of 8.8 ± 3.0 nM (R^2^ = 0.9978) (Fig. 3B). Comparison of the K_d_ values and their statistical significance by a Student’s T-test reveals that the binding trends for miR-760-3p to the 3’-UTR T100:TL duplex mimic and the full-length construct were not statistically significant from each other. In a control experiment, we titrated miR-34a-5p to the pyrC-tagged 3’-UTR T100:TL duplex mimic, and observed no quenching of fluorescent signal (S1 Fig. 3A). Additionally, pre-miR-125a was titrated to the DY547-miR-760-3p as a negative binding control for the full-length 3’-UTR, and again, we did not observe a significant quenching of the signal (S1 Fig. 3B). This data, when taken together with our previous binding native PAGE results, supports the specificity of the miR-760-3p binding interactions both to the 3’-UTR T100:TL duplex as well as to the full-length 3’-UTR in its native fold.

**Fig. 3:**
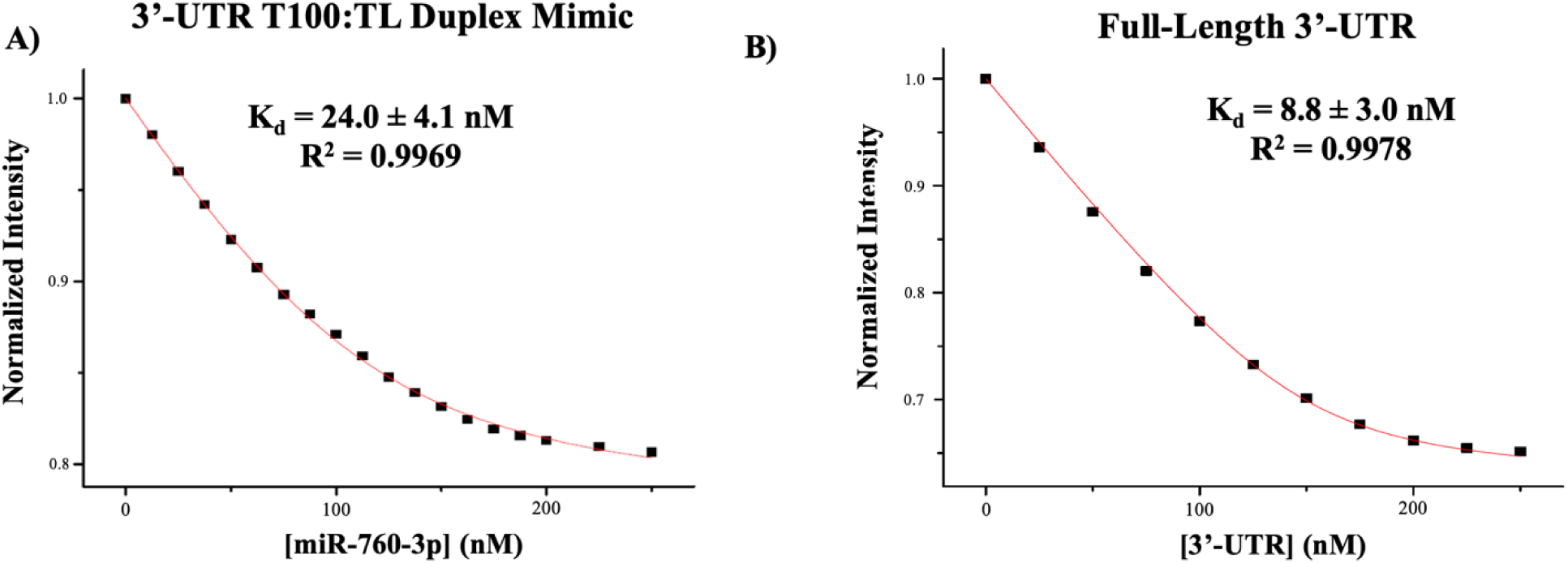
Determination of K_d_ of miR-760-3p binding to the 3’-UTR T100:TL Duplex Mimic and the full-length 3’-UTR by steady-state fluorescence spectroscopy. **(A)** K_d_ for the miR-760-3p:3’-UTR T100:TL duplex mimic was determined to be 24.0 ± 4.1 nM, and **(B)** the K_d_ for miR-760-3p:full-length 3’-UTR complex was determined to be 8.8 ± 3.0 nM. Student’s T-test (two-tailed, equal variance) of the binding curve raw data points revealed *P* (0.8318) > *ɑ* (0.01) indicating no statistically significant difference between K_d_ values.

### Host miR-34a-5p and miR-34b-5p Bind Downstream of the SARS-CoV-2 Genome 3’-UTR s2m Element

Host miR-34a-5p and miR-34b-5p both have a predicted binding site on the genome 3’-UTR downstream of the s2m element (Fig. 1, red dashed line) which is predicted to fold into a small hairpin structure (Fig. 1 red) immediately following the s2m extended structure (Fig. 1, green and yellow). (6,7) Thus, to mimic the miR-34a-5p and miR-34b-5p binding sites, we first used a sequence construct spanning the entire s2m element (Fig. 1, green), its extended lower stem (Fig. 1, yellow), and the downstream hairpin (Fig. 1, red), named here DIS-s2m extended (dimer initiation site-s2m extended). To assess binding interactions between both miR-34a-5p and miR-34b-5p to the DIS-s2m extended, we performed native PAGE experiments (Fig. 4A and 4B). It has been reported that the s2m dimerizes in the presence of Mg^2+^ through the formation of a kissing dimer, which converts to an extended duplex structure, affecting its migration in native PAGE experiments. Thus, we first analyzed the dimerization properties of the DIS-s2m extended construct in comparison to the isolated s2m (Fig. 1, green) and the DIS-s2m extended (Fig. 1, green, yellow, and red). Our results show that when incubated in the presence of increasing Mg^2+^ concentrations, the DIS-s2m extended shows a prominent dimer band (S1 Fig. 4, lanes 7-9), in contrast to the isolated s2m which as reported previously forms a mixture of monomer, kissing dimer and extended duplex conformations (S1 Fig. 4A, lanes 1-3). (6) The DIS-s2m extended dimer band could originate from a mixture of three different conformations: kissing dimer, extended duplex and dimer stabilized through a duplex formed by the extended hairpin region (Fig. 1, red) which does not involve the s2m initiated dimerization (S1 Fig. 4C). To distinguish between these possibilities, we have used a DIS-s2m construct which lacks the extended hairpin region (Fig. 1, green and yellow) and observed that this is primarily monomeric (S1 Fig. 4A, lanes 4-6). To obtain clearer data about the dimerization of these constructs, we performed the same dimerization experiments but incubated the samples in the presence of increasing Mg^2+^ concentrations for 24 hours, conditions in which we showed previously that promote formation of both kissing dimer and extended duplex conformations (S1 Fig. 4B).(6) In these conditions, the DIS-s2m remains mostly monomeric, but its dimer bands are more apparent (S1 Fig. 4B, lanes 4-6). Thus, we conclude that in addition to dimer conformations mediated by the s2m, the prominent dimer band observed for DIS-s2m extended also contains a dimer mediated through the extended hairpin, which is absent from DIS-s2m.

**Fig. 4:**
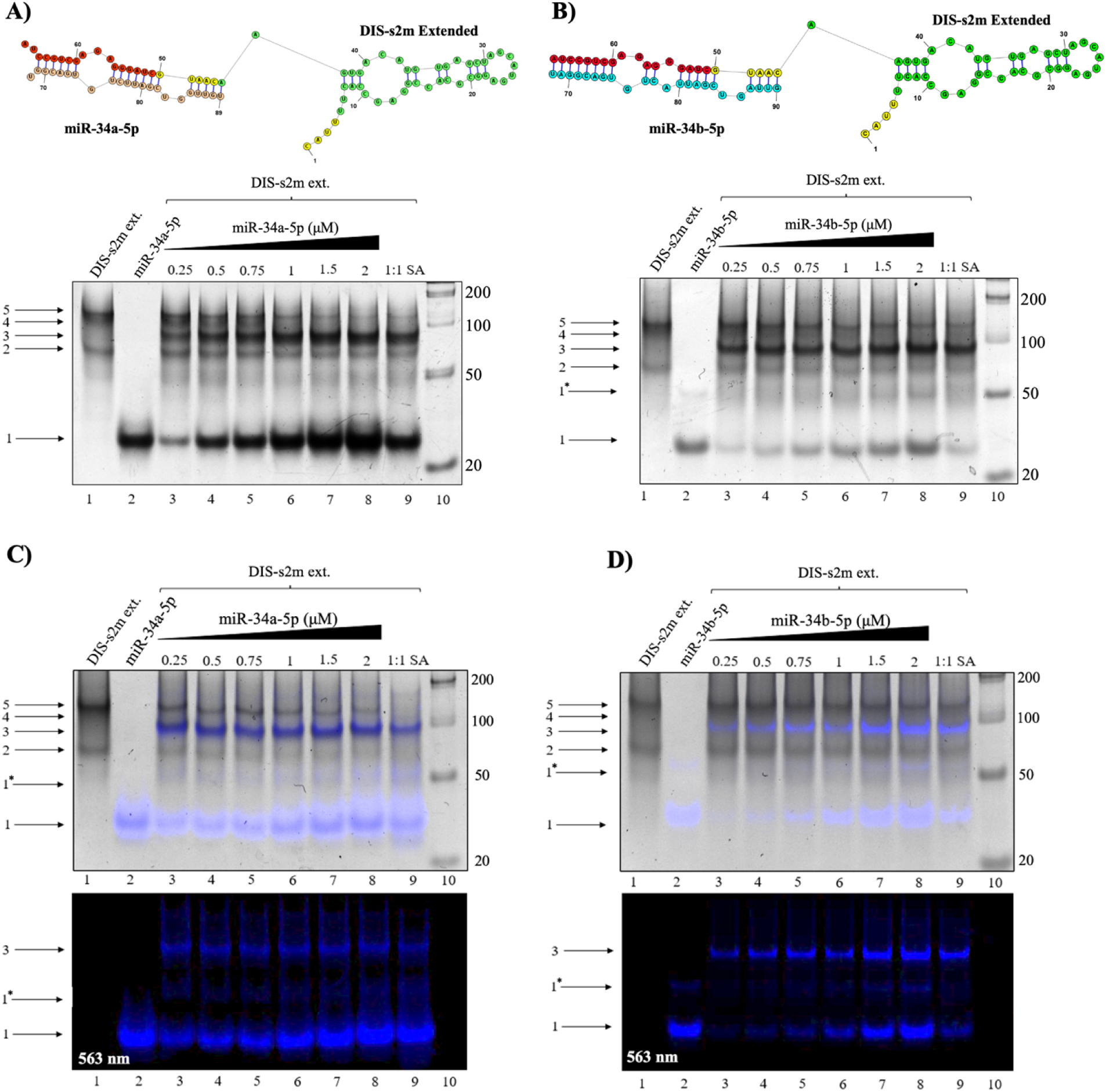
Native PAGE analysis of the binding interactions of miR-34a-5p and miR-34b-5p to the DIS-s2m extended. **(A)** Predicted structure of a 1:1 complex of DIS-s2m extended bound to miR-34a-5p (top). The DIS-s2m extended appears as a monomer (bottom, lanes 1, 3-9, arrow 2) and a mixture of dimer complexes (bottom, lanes 1, 3-9, arrows 4 and 5). Upon titration of miR-34a-5p, which exists as a monomer (bottom, lane 2, arrow 1), the DIS-s2m extended dimer complexes decrease in intensity concomitant with the appearance of a predominant binding complex (bottom, lanes 3-9, arrow 3), with migration patterns indicating a complex size of 89 nt which corresponds to the predicted structure (top). **(B)** Similarly, miR-34b-5p migrates as a monomer and dimer (bottom, lane 2, arrows 1 and 1*) is predicted to form a 1:1 complex with DIS-s2m extended (top), and titration of miR-34b-5p revealed a similar complex (bottom, lanes 3-9, arrow 3) at 90 nt which corresponds to the predicted 1:1 bound complex (top). Native PAGE analysis of the **(C)** Cy3-miR-34a-5p and **(D)** Cy3-miR-34b-5p binding complexes with the DIS-s2m extended, shown as an overlay (top), the original fluorescent image taken at 563 nm (bottom). The identity of the bands in both **(C)** and **(D)** is exactly the same as the prior experiments, with the addition of a faint band that corresponds to a miR-34a-5p dimer (bottom, lanes 2-9, arrow 1*). The presence of the fluorescent signature from the Cy3 tag in both experiments indicates the presence of miR-34a-5p and miR-34-b-5p in the previously identified bound complex bands, and confirms that the identified complexes (lanes 3-9, arrow 3 in both) contains miR-34a-5p and miR-34b-5p bound to DIS-s2m extended in a 1:1 complex, in respective experiments.

Since the nucleotides involved in miR-34a-5p and miR-34b-5p binding are located within the predicted dimerization site involving the extended hairpin, we expected that the binding of either of these miRs (Fig. 4A and 4B, bottom), will disrupt this dimer structure of the DIS-s2m extended. As discussed above, the isolated DIS s2m extended (Fig. 4A bottom, lane 1) exists in equilibrium between monomer (arrow 2) and dimer structures (arrows 4 and 5), whereas miR-34a-5p is mostly monomeric (Fig. 4A bottom, lane 2, arrow 1). Upon titration of miR-34a-5p at increasing stoichiometric ratios (Fig. 4A bottom, lanes 3-9) a new upper band appears (arrow 3), with a concomitant decrease in band intensity of both DIS-s2m extended dimer bands (lanes 3-9, arrows 4 and 5). We assign this new band (arrow 3) to the 1:1 complex of DIS-s2m extended to miR-34a-5p which is 89 nt (Fig. 4A, top). The decrease in intensity of the DIS-s2m extended dimer bands further supports the idea that the DIS-s2m extended construct forms dimers through dimerization of its 3’-hairpin tail. Given that miR-34b-5p is predicted to bind to the same site as miR-34a-5p, we performed similar native PAGE experiments and found that miR-34b-5p binds to DIS-s2m extended in the same pattern as miR-34a-5p (Fig. 4B, bottom). A concomitant decrease in DIS-s2m extended dimer bands (Fig. 4B, lanes 3-9, arrows 4 and 5) with the appearance of a new band (Fig. 4B, lanes 3-9, arrow 3) indicates the formation of a 1:1 complex of DIS-s2m extended to miR-34b-5p (Fig. 4B, top) (90 nt), similar to that of miR-34a-5p.

To confirm the assignment of the 1:1 complex as DIS-s2m extended with miR-34a-5p or miR-34b-5p complexes, we repeated the respective binding experiments using the Cy3 tagged miR-34a-5p and miR-34b-5p (excitation: 555 nm, emission: 563 nm) (Fig. 4C and 4D). The Cy3-tagged miR-34a-5p and miR-34b-5p have a migration pattern change as compared to the unlabeled miRs due to the positive charge of the Cy3 fluorophore. Using both Cy3-miR-34a-5p and Cy3-miR-34b-5p, we observed a strong fluorescent signature that overlays with the previously identified complex band (Fig. 4C and 4D, arrow 3), indicating that this complex contains each of these miRNAs. However, we noted the appearance of a lower band that migrates at about 50 nt and contains a fluorescent signature, and we assigned it to a dimer of the Cy3-miR-34a-5p (arrow 1*). These experiments, when taken with the original native PAGE experiments, indicate that both miR-34a-5p and miR-34b-5p bind to their predicted site downstream of the s2m element. In a negative control experiment, we used miR-132-3p and showed that it does not bind to the DIS-s2m extended, proving that miR34a-5p and miR-34b-5p bind it specifically (S1 Fig. 5).

Next, we used steady-state fluorescence spectroscopy to determine the K_d_ of both miR-34a-5p and miR-34b-5p complexes with DIS-s2m extended, as well as to their complexes with the full-length 3’-UTR. For binding to the DIS-s2m extended construct, we utilized a pyrC modified DIS-s2m extended, which has the pyrC located within the 3’-tail of the DIS-s2m extended (5’-AGpyrCUGC-3’, bolded in Table 1). We performed experiments monitoring the binding of Cy3 tagged miRs to the full-length 3’-UTR, similar to the DY547-miR-760-3p binding experiments. For the binding of miR-34a-5p to DIS-s2m extended, we determined a K_d_ of 11.7 ± 2.9 nM (R^2^ = 0.9988) (Fig. 5A), whereas for its binding to the full-length 3’-UTR, we determined a K_d_ of 21.3 ± 1.6 nM (R^2^ = 0.9998) (Fig. 5B). Similar experiments were performed for miR-34b-5p, where we determined a K_d_ of 6.2 ± 2.2 nM (R^2^ = 0.9975) to the DIS-s2m extended (Fig. 5C), and a K_d_ of 12.8 ± 1.3 nM (R^2^ = 0.9997) for the full-length 3’-UTR (Fig. 5D). Analysis by a two-tailed Student’s T-test with equal variance revealed that for each miRNA, there is no statistically significant difference between the binding curves to the DIS-s2m extended and the full-length 3’-UTR. Similar to the native PAGE experiments, we utilized a negative binding control of miR-132-3p for the DIS-s2m extended (S1 Fig. 6A) and pre-miR-125a for the full-length 3’-UTR (S1 Fig. 6B), and found no significant quenching of the fluorescence intensity. These findings, when paired with our native PAGE analysis, indicate that miR-34a-5p and miR-34b-5p both bind to the SARS-CoV-2 genome 3’-UTR with specificity to their predicted binding site downstream of the s2m.

**Fig. 5:**
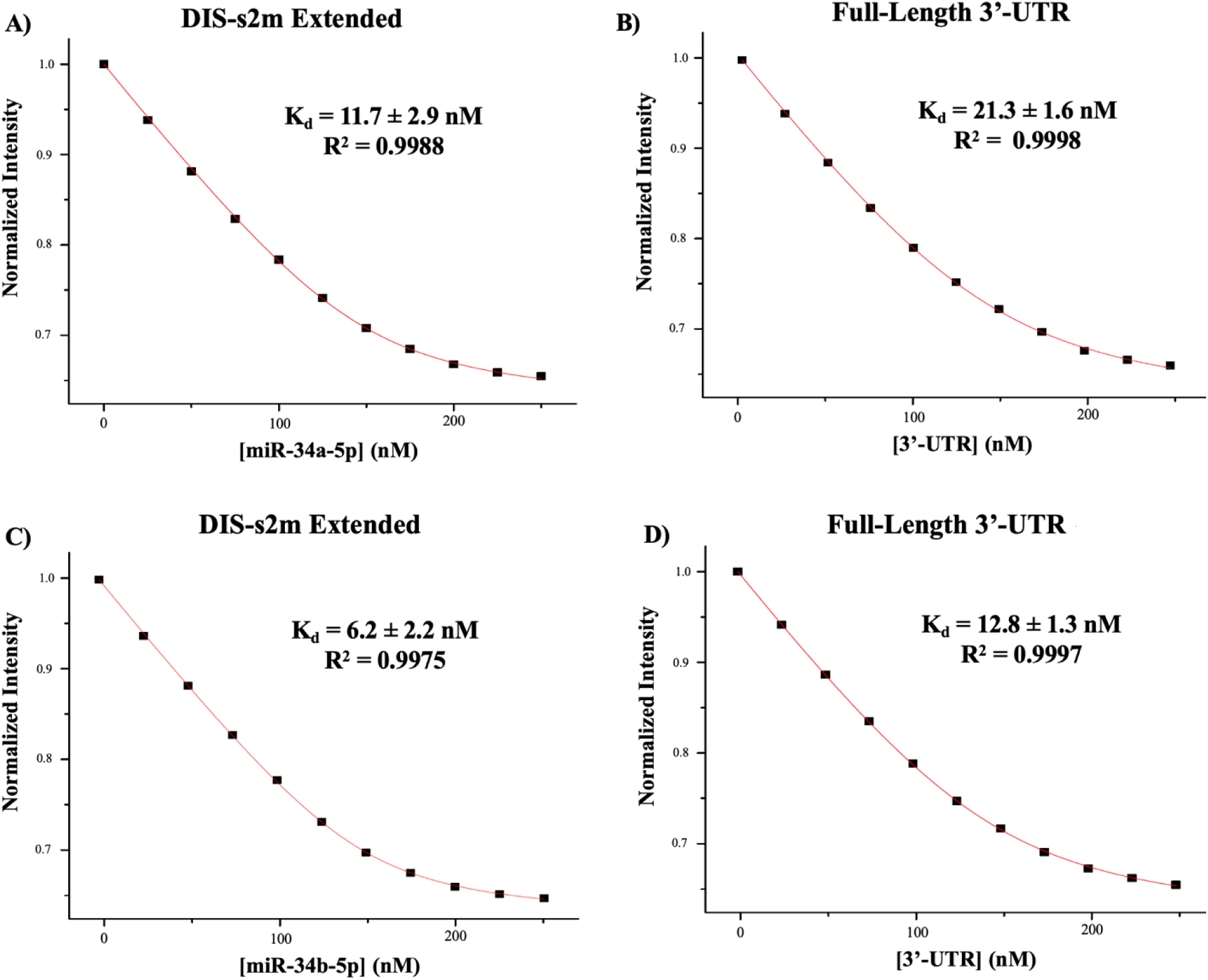
Analysis of K_d_ of miR-34a-5p and miR-34b-5p binding to the DIS-s2m extended and the full-length 3’-UTR construct by steady-state fluorescence spectroscopy. **(A)** The K_d_ for the miR-34a-5p:DIS-s2m extended complex was determined to be 11.7 ± 2.9 nM, and **(B)** K_d_ for the miR-34a-5p:full-length 3’-UTR complex was determined to be 21.3 ± 1.6 nM. Student’s T-test (two-tailed, equal variance) of the binding curve raw data points revealed *P* (0.1871) > *ɑ* (0.01) indicating no statistically significant difference between K_d_ values. **(C)** The K_d_ for the miR-34b-5p:DIS-s2m extended complex was determined to be 6.2 ± 2.2 nM, and **(D)** The K_d_ for the miR-34b-5p-full-length 3’-UTR complex was determined to be 12.8 ± 1.3 nM. Student’s T-test (two-tailed, equal variance) of the binding curve raw data points revealed *P* (0.1561) > *ɑ* (0.01), indicating no statistically significant difference between K_d_ values.

### FANA-760 and FANA-34 Bind Competitively the miR-760-3p and miR-34a-5p and miR-34b-5b Binding Sites on the SARS-CoV-2 3’-UTR

Nucleic acid analogs have been employed as therapeutic agents in a vast majority of disorders, allowing for sequence specific targeting of genomic or proteomic targets.(42,43,56,57) Of particular interest in this study, we focused on 2’-fluoro-D-arabinonucleic acid analogs (FANAs), as these oligomers have been shown to be self-delivering *in vivo* and provide effective binding to their targets within both *in vitro* and *in vivo* systems. (42–44) Thus, FANA analogs designed to bind to the miR binding sites could be potentially used as a therapeutic option as a binding inhibitor to block miR binding on the SARS-CoV-2 genome 3’-UTR, relieving the wild-type miRs to perform their normal functions in the host.

We designed a FANA analog of miR-760-3p as a perfect complement to the 3’-UTR terminus sequence, denoted as FANA-760 (Fig. 6A, top), as well as a FANA analog of miR-34a-5p and miR-34b-5p, denoted as FANA-34 (Fig. 6B, top) (Table 1, AUM Biotech). To test how these FANA oligomers interact with the 3’-UTR, we performed native PAGE experiments. We first analyzed FANA-760 binding to the 3’-UTR T100:TL duplex (Fig. 6A). FANA-760 exists primarily as a dimer (Fig. 6A, bottom, lane 3, arrow 2) with several higher molecular weight complexes being also present. We observed several binding patterns of FANA-760 to the predicted miR-760-3p binding site, including both a complex of TL:FANA-760 in a 1:1 ratio (Fig. 6A bottom, lanes 5-9, arrow 3) and a 3’-UTR TL:T100 duplex:FANA-760 complex (Fig. 6A bottom, lanes 5-9, arrow 5). Higher molecular weight complexes observed are assigned to dimers of the observed complexes, as prior PAGE experiments revealed that the TL sequence is capable of dimerizing at its 3’ end. The results that the FANA-760 oligomer is shown in PAGE to bind to both the 3’-UTR T100:TL duplex mimic and the TL sequence itself indicate that FANA-760 is capable of displacing the T100 sequence from the pre-formed 3’-UTR TL:T100 duplex, the shift in band migration indicating that the T100 oligomer is being replaced by the FANA-760 oligomer. The native PAGE analysis of FANA-34 to the DIS-s2m extended (Fig. 6B) revealed a highly similar binding pattern to that of both miR-34a-5p and miR-34b-5p to the DIS-s2m extended in which binding of FANA-34 forms a 1:1 complex with DIS-s2m extended, as seen by the appearance of a band ~ 92 nt (Fig. 6B, bottom, arrow 3). This band appears with a concomitant decrease in band intensity of both DIS-s2m extended dimer structures, which migrate as a single band (Fig. 6B, bottom, arrow 4). Additionally, we noted that two bands appeared marking a homodimer of FANA-34 (Fig. 6B, bottom, arrow 1*) and a degradation product of the DIS-s2m extended (Fig. 6B, bottom, arrow 2*).

**Fig. 6:**
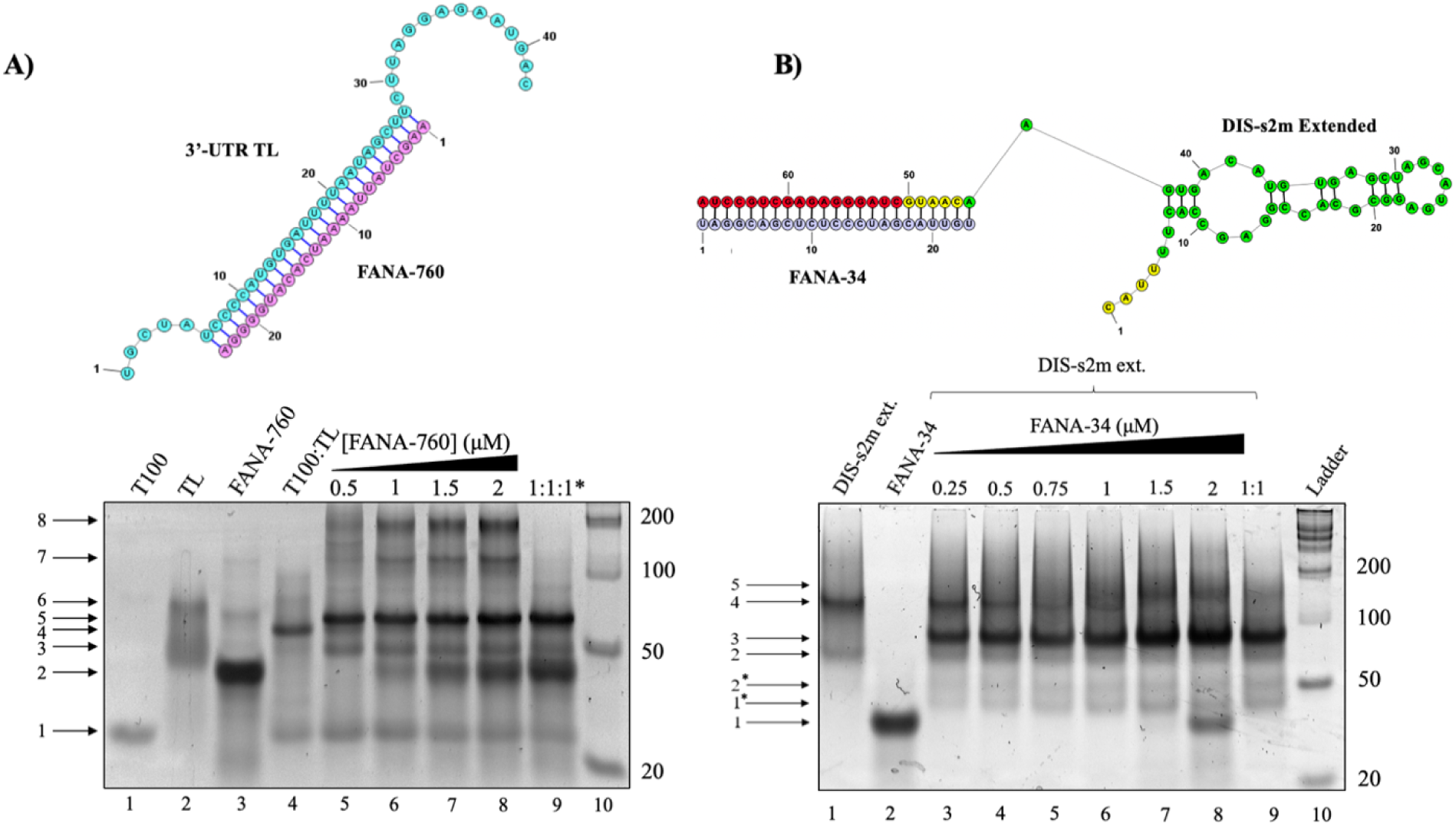
Native PAGE of the FANA-760 and FANA-34 oligonucleotides binding to the 3’-UTR duplex mimic and the DIS-s2m extended, respectively. (**A)** The FANA-760 sequence is theorized to bind to the TL sequence with competitive affinity due to its design as a perfect complement to the TL, shown by the predicted structure (top). Binding experiments by addition of FANA-760 to the 3’-UTR T100:TL duplex mimic (bottom) reveal that FANA-760 (lane 3, arrow 2) binds to the TL (lane 2, arrows 3 and 6) and forms a 1:1 complex of the TL sequence to FANA-760 (lanes 5-9, arrow 5), which is evidenced by the increasing intensity of the T100 band (lanes 5-9, arrow 1) with the addition of the FANA-760 oligomer. Additionally, we note that the FANA-760 binds to the 3’-UTR T100:TL duplex mimic (lanes 5-9, arrow 5). (**B)** The predicted structure of FANA-34 bound to the DIS-s2m extended shows perfect complementarity to the miR-34a/b-5p binding site (top). Titration of FANA-34 (lane 2, arrow 1) to the DIS-s2m extended (bottom) reveals the formation of a 1:1 complex of FANA-34 with the DIS-s2m extended (lanes 3-9, arrow 3), that appears concomitantly with a decrease in intensity of the DIS-s2m extended bands (lanes 1, 3-9, arrows 2 and 4). Notably, a higher molecular weight complex (lanes 3-9, arrow 5) is seen which is attributed to an additional molecule of FANA-34 binding to a dimer of the DIS-s2m extended. Additional bands were observed in lanes 3-9, indicated by arrows 1* and 2*, attributed to degradation products of the FANA-34 and DIS-s2m extended.

To quantitatively analyze the binding of FANA-760 and FANA-34 to the SARS-CoV-2 genome 3’-UTR, we performed steady-state fluorescence spectroscopy utilizing the pyrC-tagged 3’-UTR TL:T100 duplex and DIS-s2m extended sequences used in the wild-type miRNA steady-state fluorescence spectroscopy experiments (Fig. 7A and 7B). FANA-760 was titrated to the 3’-UTR TL:T100 duplex and FANA-34 was titrated to the DIS-s2m extended sequences, respectively, in a similar manner to prior experimentation, after which each binding curve was then fit to equation 1 to determine the K_d_ for each complex. We determined a K_d_ of 16.4 ± 1.8 nM (R^2^ = 0.9989) for FANA-760 binding to the 3’-UTR T100:TL duplex mimic (Fig. 7A) and determined a K_d_ of 8.7 ± 1.0 nM (R^2^ = 0.9997) for FANA-34 binding to the DIS-s2m extended (Fig. 7B), noting that the FANA oligomers have competitive K_d_ values to the wild-type miRs. Thus, we performed competition assays for further characterization of the FANAs as binding inhibitors, calculating the IC50 and K_I_ for each FANA. In these competition assays, we incubated the Cy3- or DY547-tagged miRNAs with the full-length 3’-UTR and titrated the respective FANA, observing an increase in the fluorescence intensity as the miRNAs are competed off by the respective FANA. The same experiments were also repeated by titrating an unlabeled miR-760-3p or miR-34a/b-5p, monitoring the displacement of the fluorescent tagged miRs can be monitored as was done for the FANAs to allow the comparison of the IC50 values for the FANA and the respective miR it displaced. The fluorescence intensities in all competition assays were normalized to the first data point prior to titration of the unlabeled ligand (FANA or miRNA) and fit with equation 2 to determine IC50 (Fig. 7C and 7D). The IC50 values were then analyzed using equation 3 to determine the K_I_ values. For FANA-760, we determined an IC50 of 107.7 ± 12.7 nM, and for comparison, miR-760-3p was found to have an IC50 of 124.1 ± 9.8 nM (Fig. 7C). The K_I_ of FANA-760 was calculated to be 6.0 ± 0.2 nM. Analysis of FANA-34 revealed an IC50 of 56.1 ± 2.1 nM, while miR-34a-5p was determined to have an IC50 of 117.4 ± 7.8 nM and miR-34b-5p was determined to have an IC50 of 116.5 ± 6.7 nM (Fig. 7D). From this information, a K_I_ of 7.0 ± 1.9 was calculated for FANA-34 to the full-length 3’-UTR construct (Fig. 7D). As a negative control for the competition assay, we titrated miR-132-3p in place of the FANA-760 and FANA-34, and found no increase of the fluorescence intensity (S1 Fig. 7), supporting the ability of the FANA oligomers to compete with the wild-type miRNAs for binding to the full-length SARS-CoV-2 genome 3’-UTR and confirming that the interactions are sequence specific.

**Fig. 7:**
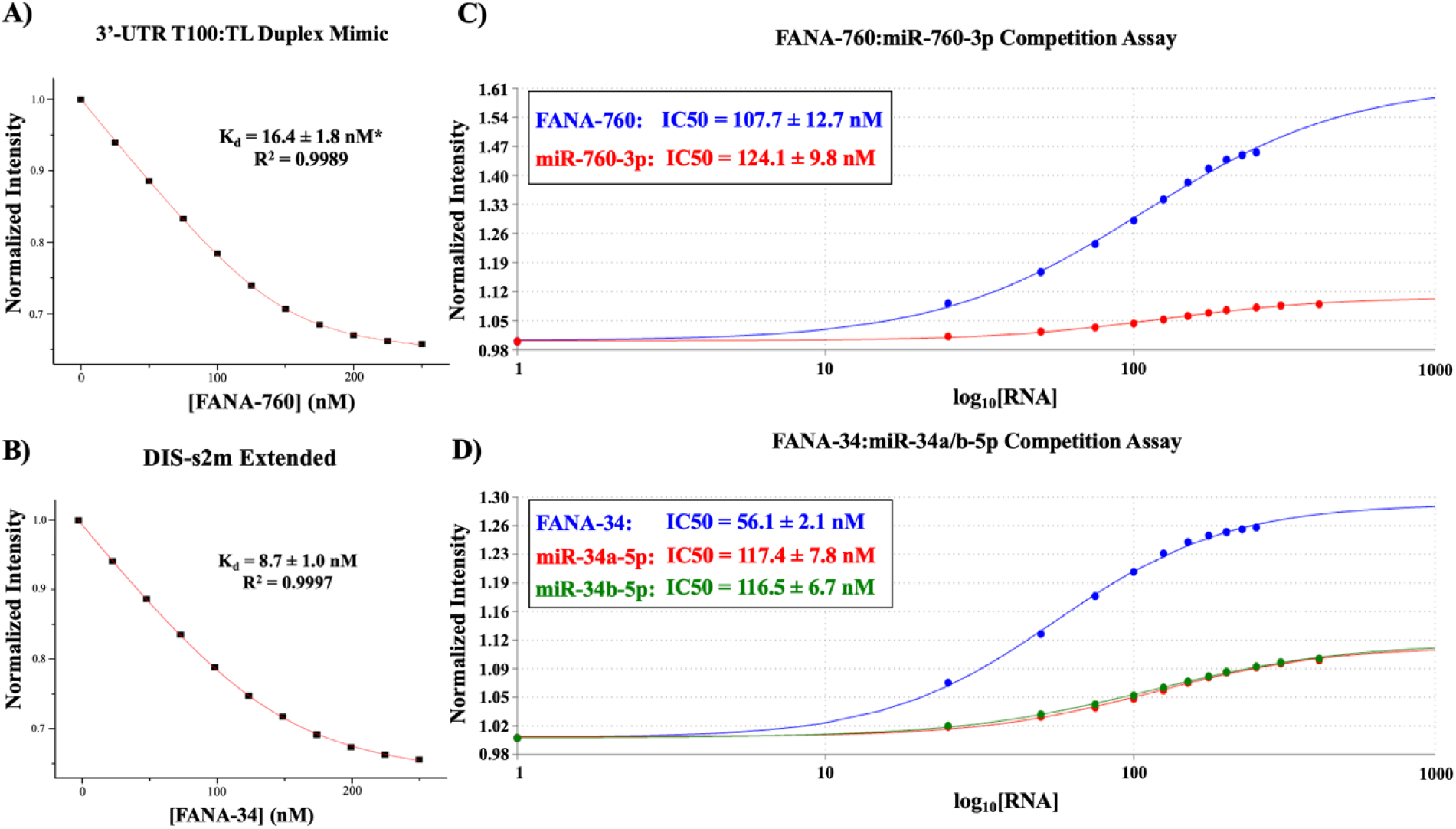
Determination of FANA-760 and FANA-34 binding interactions to the SARS-CoV-2 3’-UTR by steady-state fluorescence spectroscopy. **(A)** The K_d_ for FANA-760 to the 3’-UTR T100:TL duplex complex was determined to be 16.4 ± 1.8 nM. **(B)** The K_d_ value for FANA-34:DIS-s2m extended was determined to be 8.7 ± 1.0 nM. **(C)** Competition assays allowed the determination of IC50 values of 107.7 ± 12.7 nM for FANA-760 and 124.1 ± 9.8 nM for miR-760-3p binding to the full-length 3’-UTR, and **(D)** IC50 values of 56.1 ± 2.1 nM for FANA-34, 117.4 ± 7.8 nM for miR-34a-5p, and 116.5 ± 6.7 nM for miR-34b-5p binding to the full-length 3’-UTR.

## Discussion

In this work, we demonstrate that miR-760-3p, miR-34a-5p, and miR-34b-5p bind specifically and with high affinity to the SARS-CoV-2 genome 3’-UTR. In several other viruses, miR “sponging,” the process of binding miRs to viral nucleic acids and/or proteins, has been directly correlated with virulence and malignancy.(55,58,59) While our work is performed *in vitro*, our experimentation is performed in conditions that mimic the native fold of the SARS-CoV-2 genome 3’-UTR as well as the physiological concentration of Mg^2+^. For miR-760-3p binding, native PAGE reveals the formation of a 1:1 complex between the 3’-UTR T100:TL duplex mimic and miR-760-3p. Given that the 3’-UTR T100:TL duplex mimics the native fold of the 3’-UTR terminus *in vivo,* this finding suggests that host miR-760-3p is capable of binding to the SARS-CoV-2 genome 3’-UTR without disrupting its native fold.(5) Further, we report a K_d_ for miR-760-3p binding to the 3’-UTR T100:TL duplex mimic that is not statistically different from its binding to the full-length 3’-UTR, which increases the relevancy of this interaction *in vivo* as the 3’-UTR construct was expressed and purified under native conditions, mimicking the native 3’-UTR fold in the full length genome. Thus, our results support the prediction that miR-760-3p binds the SARS-CoV-2 genome *in vivo*, and suggest a potential mechanism of miR-760-3p hijacking by SARS-CoV-2 in infected cells with several implications in the viral life cycle.

For the binding interactions of miR-34a-5p and miR-34b-5p, our data indicates that both miRNAs bind to the DIS-s2m extended in a similar binding pattern, with each miRNA exhibiting a strong binding affinity. All these miRNAs bind to their predicted targets *in vitro*, which could imply that interactions of the these predicted viral binding sites with other portions of the viral genome could interfere with binding of these miRNAs. It is known that the SARS-CoV-2 viral genome is highly dynamic, and thus, long-range or short-range interactions that are formed between the 3’-UTR and other regions/copies of the viral genome could potentially influence binding interactions of miR-760-3p, miR-34a-5p, and miR-34b-5p.(4,6,60) Additionally, with the finding that both miR-34a-5p and miR-34b-5p could bind to the coding regions of the viral M and S proteins, respectively, we propose that the binding of these miRNAs to the 3’-UTR could serve as a molecular “decoy” by providing an alternative site for targeting copies of the viral genomic and subgenomic RNAs which contain the genome 3’-UTR.(33) MiR-760-3p has also been identified to regulate the ACE2 receptor mRNA 3’-UTR, and such, hijacking of host miR-760-3p could lead to an upregulation of ACE2, ultimately favoring viral entry into infected cells.(26) Taken together, the hijacking of miR-760-3p, miR-34a-5p, and miR-34b-5p could benefit the virus by reducing the targeting of viral structural proteins by host miR-34a-5p and miR-34b-5p as employed by the immune response, and increasing viral entry through increased expression of ACE2. These factors combined would aid in SARS-CoV-2 viral fitness, and the binding mechanisms we characterize would facilitate these interactions.

It is known that in CRS, a large portion of proinflammatory biomolecules, like IL-6, IL-6R, and PGRN, are dysregulated. Given the previously reported regulation of these biomolecules by miR-760-3p, miR-34a-5p, and miR-34b-5p, binding interactions between these miRs and SARS-CoV-2 may trigger the dysregulation of the immune response through a loss of translational repression.(20,21,21) Continuous positive feedback cycles, like those observed in CRS, could be amplified given increased binding of these miRs in parallel with increased presence of SARS-CoV-2 RNAs which harbor the 3’-UTR. Thus, more severe infections that result in higher viral titers may be attributed to increased disease severity and CRS through the hijacking of these miRs, as more miRs may be “sponged” by the virus. Clinical data on SARS-CoV-2 revealed that large viral titers, upregulation of immune activity, and an apparent loss of translational inhibition of core cytokines have been attributed to CRS development and overall severity, highlighting a potential implication of miR hijacking by SARS-CoV-2 and increasing the relevancy of these interactions *in vivo*.(36,38,39). Although the recent variants of SARS-CoV-2 are less severe than the original Wuhan strain, severe symptoms associated with SARS-CoV-2 infection still pose a danger to those that are immunosuppressed or compromised, and highlights a need for therapeutic intervention.(37,38,61) Our implementation of the FANA-760 and FANA-34 oligomers allows for a direct targeting of the miR hijacking interactions, as we have shown by our *in vitro* analysis that these oligomers are capable of competing with the miR binding interactions with SARS-CoV-2.

Our characterization of both FANA-760 and FANA-34 concurs with our analysis of the wild-type miRs, and emphasizes that these FANA oligomers bind with specificity to the SARS-CoV-2 genome 3’-UTR at their targeted sites. Our fluorescence spectroscopy data shows that both FANA-760 and FANA-34 are capable of binding with high affinity to the 3’-UTR at their respective predicted binding sites, and are also capable of competing with the wild-type miRs for the full-length 3’-UTR. FANAs have been applied to other *in vivo* systems, allowing for targeted interactions with sequence specificity.(42–44) FANAs have been shown to be self-delivering, which allows for the FANAs to act with limited accessories related to delivery to target cells. While we do not present *in vivo* work, our competition assays reveal that the FANAs have competitive IC50 values to those of the wild-type miRs to the full-length SARS-CoV-2 3’-UTR. These IC50 values, along with the respective K_I_ for the FANAs, are physiologically relevant and plausible for implementation in live organisms.(62,63) We propose that the FANA-760 and FANA-34 oligomers, both of which are sequence specific to SARS-CoV-2, could be used as direct binding inhibitors to SARS-CoV-2, inhibiting miR hijacking to the 3’-UTR and relieving miR-760-3p, miR-34a-5p, and miR-34b-5p to perform their natural functions and restore regulation to the immune response.

## Conclusions

In this study, we identify and characterize *in vitro* the binding interactions of novel targets of viral hijacking by the SARS-CoV-2 virus in miR-760-3p, miR-34a-5p, and miR-34b-5p, and we implicate the potential role of these interactions in viral life cycle. We also propose that the FANA analogs of these miRs, FANA-760 and FANA-34, can be utilized in the mitigation of these binding interactions though intervention as binding inhibitors. We emphasize that the IC50 and K_I_ values determined in this work for these FANAs are consistent with literature on biologically relevant interactions of drugs with their targets.(62,63) Altogether, we anticipate that the binding interactions between miR-760-3p, miR-34a-5p, and miR-34b-5p are relevant in severe COVID-19, as well as in the host immune response to infection, and that implementation of FANAs like FANA-760 and FANA-34 could provide a novel route to antiviral therapy through restoration of host immune function.

## Author Contributions

MRM conceived of the presented idea and supervised the work. CJF performed the experiments and data analysis, based on preliminary work by CLC. CJF wrote the initial manuscript and worked with CLC and MRM for preparation of the revised manuscript prior to submission.

## Acknowledgements

This work was supported by the National Science Foundation Division of Chemistry Rapid Response Research (RAPID) CHE 2029124, and National Institutes of Health 2R15GM127307-05.

## Supporting Information Figure Legends

**S1 Fig. 1: Native PAGE analysis of the miR-760-3p binding interactions to the 3’-UTR duplex mimic with the TL mutant sequence.** To confirm the specific interactions of miR-760-3p to the exposed bulge formed in the 3’-UTR T100:TL duplex mimic, we mutated the exposed bulge on the TL sequence and repeated miR-760-3p binding experiments. The T100 monomer (lane 1, arrow 1) and TL mutant sequence (lane 2, arrows 2 and 5) were slow annealed similar to prior experiments, forming the 3’-UTR T100:TL duplex mimic (lane 4, arrow 4). Upon titration of miR-760-3p, we observe no binding to the 3’-UTR T100:TL duplex mimic. Several smaller molecular weight bands are observed (lanes 2-9, bracket 2*), and we attribute these bands to degradation products of the TL mutant and miR-760-3p sequences.

**S1 Fig. 2: Native PAGE of miR-34a-5p and miR-1307-3p binding to the 3’-UTR T100:TL duplex mimic.** To establish sequence specificity of the 3’-UTR T100:TL duplex mimic for miR-760-3p, two microRNAs, miR-34a-5p (lanes 3, 6, and 7, arrow 2) and miR-1307-3p (lanes 4, 8, and 9, arrow 4) were utilized as controls. The free T100 (lane 1, arrow 1) and TL (lane 2, arrows 3 and 6) sequences were run to confirm the identity of the free RNAs in later samples. Preformation of the 3’-UTR T100:TL duplex mimic (lanes 4-9, arrow 5), followed by titration of either miR-34a-5p or miR-1307-3p, did not result in the appearance of any higher molecular weight complex bands. No change in intensity of the 3’-UTR T100:TL duplex band (lanes 4-9, arrow 5) was observed, supporting no binding between the 3’-UTR T100:TL duplex mimic and the control miRs.

**S1 Fig. 3: Steady-state fluorescence spectroscopy of the negative binding control for the 3’-UTR T100:TL duplex mimic. (A)** Titration of miR-34a-5p, which was previously shown not to bind in native PAGE, was performed as a negative binding control for the miR-760-3p binding experiments. Upon incremental addition of miR-34a-5p, no significant change was observed in the fluorescence intensity of the pyrC-tagged 3’-UTR T100:TL duplex mimic. **(B)** For the full-length 3’-UTR experiments, pre-miR-125a was used in place of the 3’-UTR and titrated to DY547-miR-760-3p as a negative control. No significant decrease in fluorescence intensity was observed, indicating specificity of the 3’-UTR for miR-760-3p.

**S1 Fig. 4: Comparison of the isolated s2m, DIS-s2m, and DIS-s2m extended dimerization by native PAGE. (A)** While the isolated s2m shows the formation both kissing dimer and extended duplex dimers (lanes 1-3, arrows 4 and 5), extension of the motif to include its 4 lower base pairs (DIS-s2m) limits dimer formation. A construct containing the DIS-s2m and a 3’-tail, which contains the miR-34a/b-5p binding sites (DIS-s2m extended) forms a dimer that is formed through the dimerization of this tail (lanes 7-9, arrow 7). We also observe a kissing dimer (lanes 7-9, arrow 6) that is consistent with the isolated s2m dimerization pattern, explaining the uppermost complex bands of the free DIS-s2m extended. **(B)** Incubation at 37°C for 24 hours revealed greater dimer formation for all three constructs, indicating that the dimer conformations are thermodynamically and kinetically favored. **(C)** Structures of the DIS-s2m extended as a kissing dimer (top), a duplex mediated by the kissing dimer (middle), and a dimer mediated by the 3’-hairpin (bottom).

**S1 Fig. 5: Native PAGE analysis of the negative binding control of miR-132-3p to the DIS-s2m extended.** Titration of miR-132-3p as a specificity control to the DIS-s2m extended revealed no binding interactions, as no apparent change in the intensity of the DIS-s2m extended is observed (lanes 2-9, arrows 2 and 3) upon incremental addition of miR-132-3p.

**S1 Fig. 6: Steady-state fluorescence spectroscopy analysis of the negative binding control of miR-132-3p to the DIS-s2m Extended. (A)** The titration of miR-132-3p incrementally to the DIS-s2m extended revealed no change in fluorescent intensity. **(B)** Titration of pre-miR-125a to the Cy3-tagged miR-34a/b-5p oligomers revealed no change in fluorescent intensity, consistent with prior controls in demonstrating sequence specificity of these interactions for the target miRNAs.

**S1 Fig. 7: MiR-132-3p as a control for the FANA-miRNA competition experiments.** To demonstrate the specific activity of FANA-760 and FANA-34 in their competition with the wild-type microRNAs, we titrated miR-132-3p in place of the FANAs, repeating the competition experiment. The fluorescent intensities were normalized to the initial intensity, in which there is 150 nM of miRNA to 250 nM of 3’-UTR. Where previously the FANAs demonstrate a gain in fluorescent intensity upon titration, we observe no significant regain fluorescent intensity upon addition of miR-132-3p.

## Equations

**Equation 1:** The model equation used for fitting of steady-state fluorescence experiments to determine the K_d_ of the microRNAs or FANAs to the SARS-CoV-2 3’-UTR T100:TL duplex, DIS-s2m extended, or full-length 3’-UTR. I_B_/I_F_ represents the ratio of fluorescence intensities of the bound (I_B_) and free (I_F_) RNA states. [X]_t_ and [Y]_t_ represent the concentration of the titrant miRNA/RNA and the concentration of the fluorescently-tagged RNA (pyrC, DY547, Cy3-tagged RNAs), respectively.

**Equation 2:** The model equation used for fitting of the competition experiments to the full-length 3’-UTR. F_o_ represents the fluorescence intensity at any specific concentration of unlabeled titrant, in this case either FANA or unlabeled miR, denoted as [S]_o_. F_max_ and F_min_ represent the maximum and minimum fluorescence intensities of the experiment, respectively.

**Equation 3:** The model equation used for calculation of K_I_ from the determined IC50 values from equation 2. [L] represents the concentration of fluorescent ligand, in this case, DY547/Cy3-tagged miR. The K_d_ used in this equation is the experimentally derived K_d_ for each FANA or miR.

